# Assessing the statistical reporting quality in high-impact factor urology journals

**DOI:** 10.1101/2020.03.19.998765

**Authors:** Shuangyang Dai, Hong Xu, Beibei Li, Jingao Zhang, Xiaobin Zhou

**Affiliations:** Department of Epidemiology and Health Statistics, School of public health, Qingdao University, No. 38 Dengzhou Road, Qingdao, Shandong Province, 266021, People’s Republic of China; Department of orthodontics, the affiliated hospital of Qingdao University. School of Stomatology, Qingdao University

**Keywords:** Urology, Observational studies, Statistical methods, Reporting quality

## Abstract

**Backgrounds:** Observational studies plays an important role in urology studies, But few studies have paid attention to the statistical reporting quality of observational studies. The purpose of this study was to investigate the frequency and evaluate the reporting quality of statistical methods of the published observational studies in urology.

**Methods:** The five urology journals were selected according to the 5-year impact factor. A systematic literature search was performed in PubMed for relevant articles. The quality of statistical reporting was assessed according to assessment criteria.

**Results:** A total of 193 articles were included in this study. The mean statistical reporting score of included articles was 0.42 (SD=0.15), accounting for 42% of total score. The items that must be reported with a reporting rate more than 50% were: alpha level (n=122, 65.2%), confidence intervals (n=134, 69.4%), name of statistical package (n=158, 84.5%) and exact *P*-values (n=161, 86.1%). The items with a reporting rate less than 50% were: outliers (n=2, 1.0%) and sample size (n=13, 6.7%). For multivariable regression models (liner, logistic and Cox), variables coding (n=27, 40.7%), validation analysis of assumptions (n=58, 40.3%), interaction test (n=43, 30.0%), collinearity diagnostics (n=5, 3.5%) and goodness of fit test (n=6, 5.9%) were reported. Number of authors more than 7(OR=2.06, 95%CI=1.04-4.08) and participation of statistician or epidemiologist (OR=1.73, 95%CI=1.18-3.39) were associated with the superior reporting quality.

**Conclusion:** The statistical reporting quality of published observational studies in 5 high-impact factor urological journals was alarming. We encourage researchers to collaborate with statistician or epidemiologist. The authors, reviewers and editors should increase their knowledge of statistical methods, especially new and complex methods.

## Introduction

Nowadays, more and more medical researchers notice the phenomenon that the main results of some research cannot be reproduced (1, 2), and improper use of statistical methods or inadequate statistical reporting may be the important reasons for this phenomenon. The low-quality statistical reporting may not make full use of research results, resulting in a waste of valuable information and varying degrees of bias. In addition, it may also make editors and readers unable to measure the reliability of research, and make readers misinterpret the results of the study, thus leading to errors in the secondary studies (3).

Truthly, the poor statistical reporting problem is long-standing, but few people pay attention to it. In 1966, the first paper on the quality of medical literature statistical report was published (4), and then dozens of similar studies were published (5). These studies found that the application of statistical methods is becoming more and more complicated, but the problem of insufficient statistical reporting has always existed. To make matters worse, the literature evaluated by these studies was from influential general medical and professional journals.

In 2015, T.A. Lang *et al.* published the “Statistical Analyses and Methods in the Published Literature’’ (SAMPL) guidelines to improve the quality of basic statistical reporting (6). The principle of SAMPL is that “authors should describe statistical methods with sufficient detail to enable readers in the professional domain to access raw data to verify the results of the report”. In 2017, Pentti Nieminen *et al.* made the SIMA (Statistical Intensity of Medical Articles) tool and assessed the statistical intensity in the high impact factor respiratory journal’s articles, and found that approximately one third of the respiratory papers provided incomplete description of their statistical reports (7, 8). Even though the SAMPL guidelines and SIMA broaden the standards for the scrutiny of statistical methods, there is still a void in requiring or assessing the reporting of statistical methods in observational studies.

Therefore, we carried out this study to describe the frequency and trends of statistical methods used in high-impact urology journals, and evaluate the reporting quality of statistical methods. We hope this study can identify the major quality deficiencies in the statistical reporting of urological observational studies, and promote the improvement of statistical reporting quality.

## Material and Methods

### Journals selection and search strategy

Top five urology journals were selected according to 5-year impact factor (excluding review journals), including European Urology (IF=17.581), Journal of Urology (IF=5.157), BJU International (IF=4.688), Prostate Cancer and Prostatic Diseases (IF=4.099), and Prostate (IF=3.820). The relevant studies from January 1, 2009 to September 30, 2019 were searched in the PubMed. The search strategy was shown in Figure. 1.

**Figure 1.**
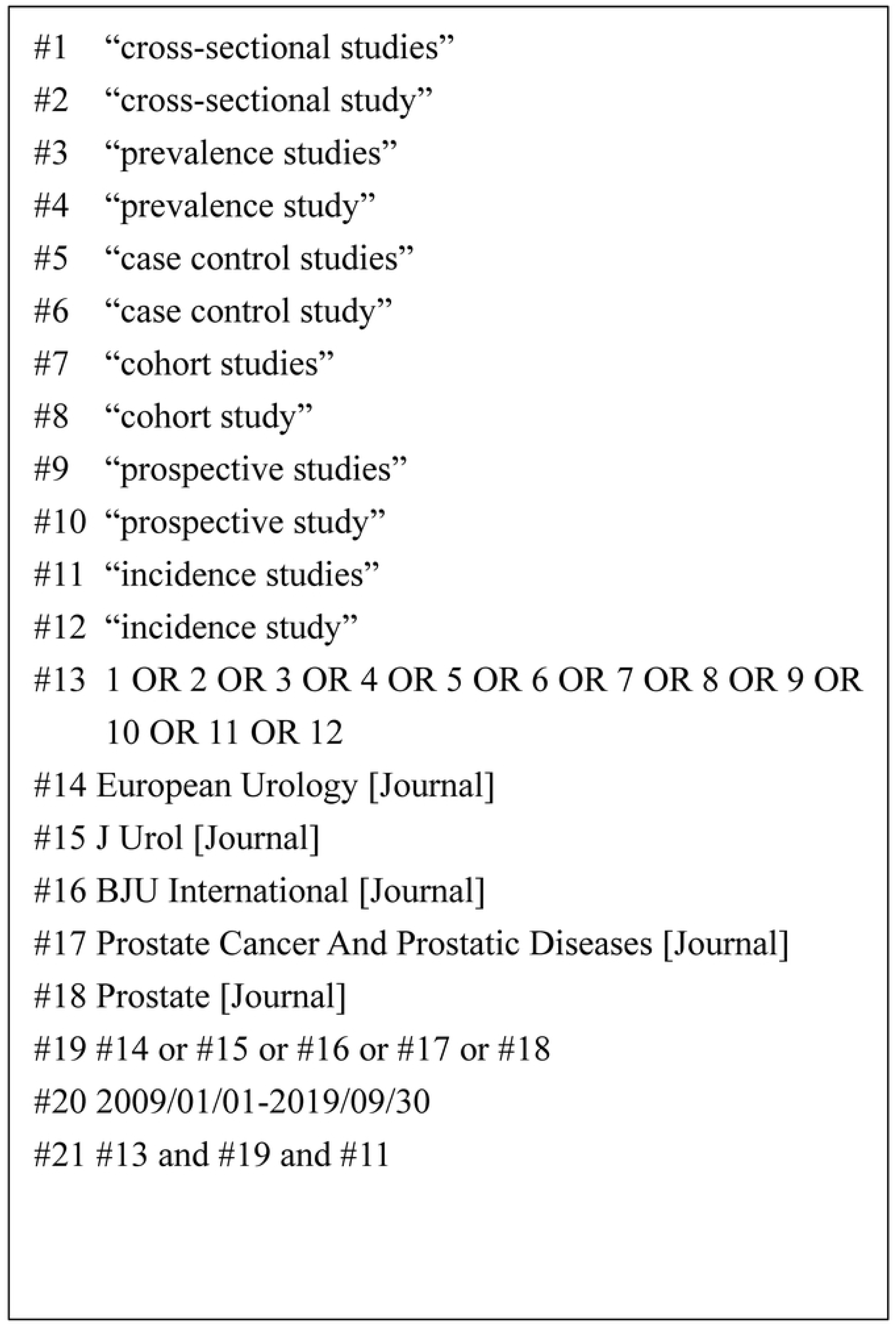
Search strategy for PubMed.

### Articles selection

Articles that met the following criteria were selected: (1) original articles; (2) observational studies, including cross-sectional studies, case-control studies, and cohort studies; (3) studies on humans, including both adults and children. The exclusion criteria were as follows: (1) review articles and case reports, (2) quasi-randomized trial, randomized controlled trials, and other interventional studies, (3) unpublished data and published abstracts only.

The articles retrieved were preliminary reviewed according to titles and abstracts by two investigators independently. Any disagreement was resolved by consulting with senior authors. After the initial screening, two investigators retrieved the full texts of relevant researches and determined the final list based.

### Frequency and trends of statistical methods used in the included articles

Two investigators extracted data independently from the included articles. Statistical methods used in the article and general characteristics were extracted, including name of journal, publication time, origin of corresponding author, type of study, number of authors, impact factor, funding support, the affiliation of corresponding author, international collaborative authorship, and participation of statistician or epidemiologist.

A standardized evaluation form (see Appendix 1) based on Emerson *et al.*(9) was used to record the frequency and trends of statistical methods for observational studies published in the selected urology journals. The original study divided statistical methods into 21 categories, 20 categories with the exception of benefit analysis have been incorporated into this checklist with slightly modified. The items containing the following information have been added to this checklist: multiple comparisons, repeated measurement data analysis, consistent measurement, Bayesian analysis.

### Statistical reporting quality of the included articles

Two reviewers applied the assessment criteria (see Appendix 2) independently to appraise the statistics reporting quality of included studies. This checklist was established based on the SAMPL guidelines (6) and other previously published studies (10-12), and the items were modified to be listed in a simple and readable manner. All of the logistic regression assessment items droved from Zhang’s research (13), and Cox regression items were from Zhu’s research (14). The checklist consists of 7 items (marked *) that must be reported and 39 items that are subject to selective reporting based on the statistical methods used. The proportions of items that were adequately reported in the statistical methods used were calculated as statistical reporting quality score for each article. The total score of statistical reporting score of each article was 100% (1). The evaluations that two investigators argued over were resolved by discussion with the senior author.

### Statistics analysis

Continuous variables with normal distribution were represented by mean ± standard deviation (SD), non-normal variables by medians (interquartile range), and categorical variables by frequency (percentage). The comparisons of means between the two groups were performed by independent Student’s t-test, and multi-group by one-way analysis of variance (ANOVA). The Mann-Whitney U test and Kruskal-Wallis test were used to analyze variables for non-normal variables. According to a cutoff value of the 75 percentile of the statistical reporting quality score, the articles were divided into high and low quality groups. Univariate and multivariate logistic regression analyses were performed to identify the factors affecting statistical reporting quality. The variables with a *P* ≤0.05 for univariate logistic analyses were included in the multivariable logistic regression model. All the statistical analyses were performed using SPSS version 21.0. All significance tests were two-sided, and *P* ≤ 0.05 was considered as significant.

## Results

### Search results

The initial search of databases confirmed 8,605 potentially relevant articles without duplication. After the screening of titles and abstracts, 8,326 articles were excluded. In total, 279 full-text articles were further reviewed, with 84 articles were subsequently eliminated for various reasons. Finally, 193 relevant articles were included. The process of literature retrieval was shown in the flow diagram (Figure.2).

**Figure 2.**
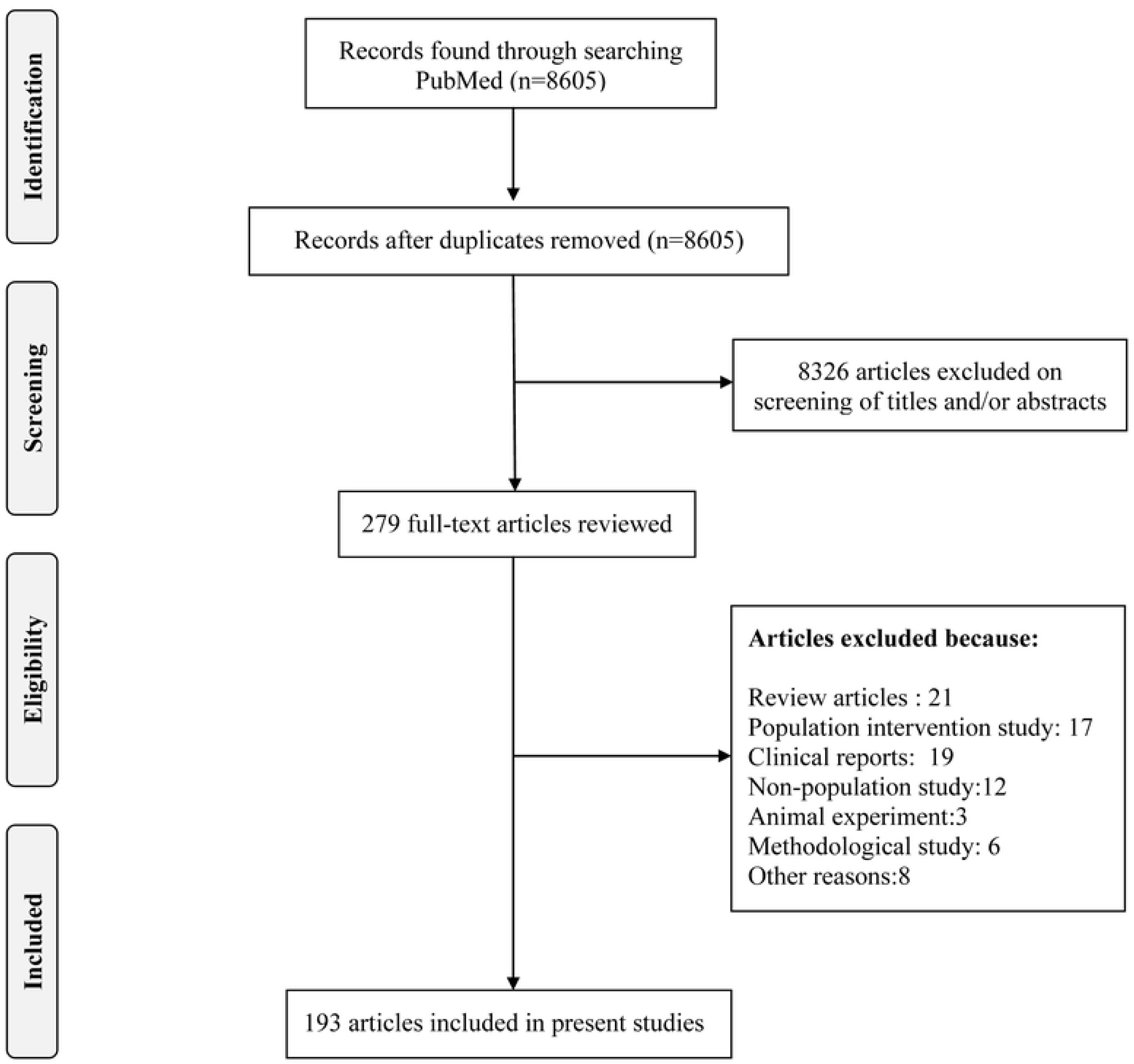
Flow diagram of the screening articles.

### General characteristics of the included articles

Of the 193 articles, 58 (30.05%) were cross-sectional studies, 56 (29.02%) were case-control studies, and 79 (40.93%) were cohort studies. Characteristics of included articles were shown in Table.1.

**Table 1.**
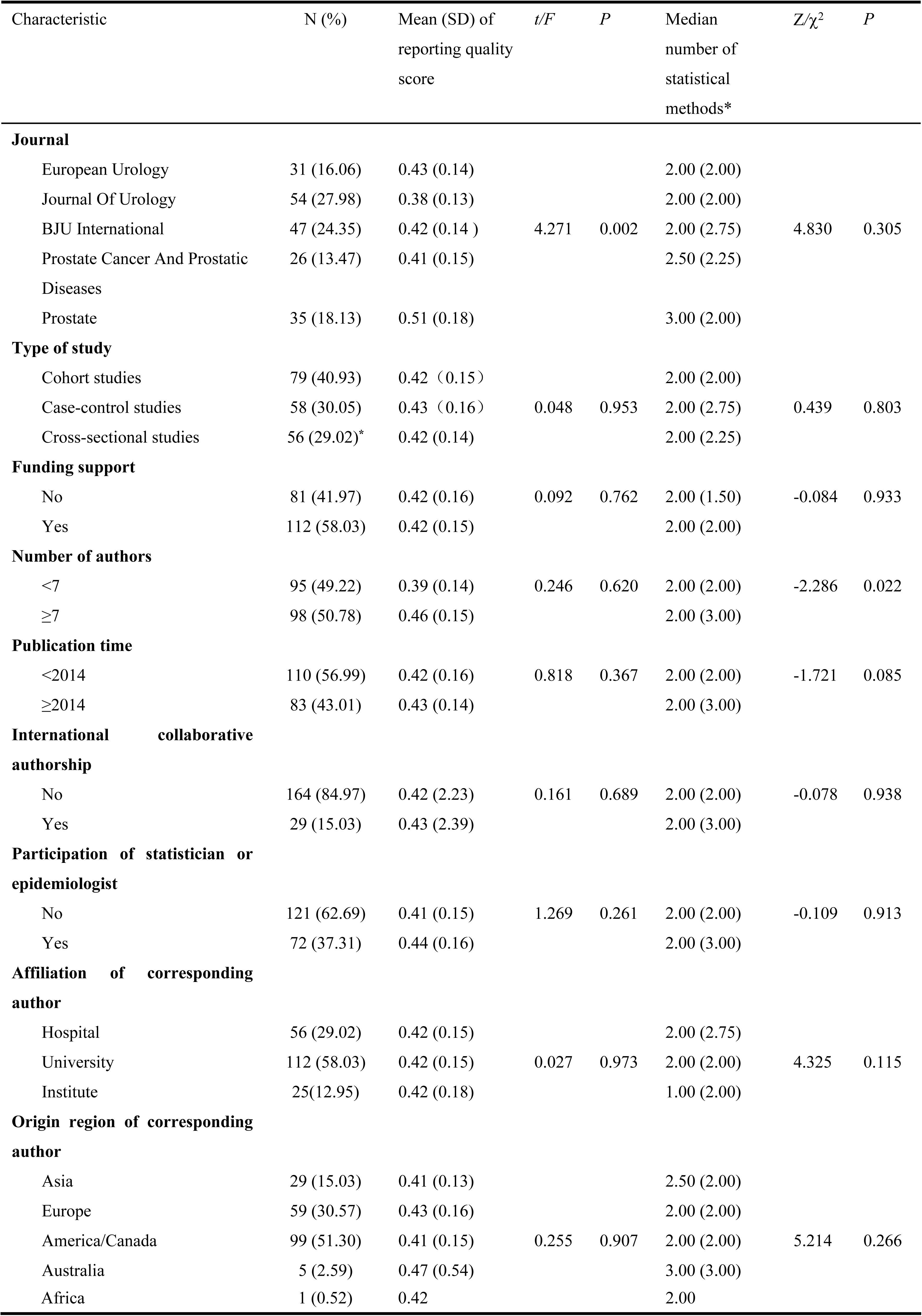
The general characteristics of included studies.

### The frequency of statistical methods applied in the included articles

The frequency of statistical methods in included articles was shown in Table.2. 6 (3.1%) articles didn’t use statistical methods or used descriptive statistics only. 36 (18.7%) articles only used simple methods such as t-test and chi-square test, 54 (28.0%) articles only adopted multivariable analyses like logistic regression and cox regression, 97 articles (50.3%) both employed simple and multivariable analyses. Logistic regression (n=93, 48.2%) and Chi-square test (n=90, 46.6%) were the most frequently used statistical methods in included studies. 7 articles (3.6%) used propensity score method. None of the articles used Bayesian methods, artificial neural networks and machine learning. The number of statistical methods in articles with 7 or more authors was more than others. The trend of the percentages of the different statistical methods used over time was shown in Figure.3.

**Table 2.** Common statistical methods in medical studies.

|  |  |  |
| --- | --- | --- |
| Descriptive statistics |  | 190 (98.4%) |
|  | Means (standard deviation) | 165 (85.5%) |
|  | Median (interpercentile range) | 39 (20.2%) |
|  | Proportion | 122 (63.2%) |
|  | Rate (e.g. incidence rate, survival rate) | 7 (3.6%) |
|  | Ratio (e.g. odds ratios, relative risk) | 11 (5.7%) |
| Ancillary analyses |  | 47 (24.4%) |
|  | Variable transforms | 19 (9.8%) |
|  | Variable constructs | 7 (3.6%) |
|  | Standardizing | 6 (3.1 %) |
|  | Matching | 0 (0) |
|  | Propensity score methods | 1 (0.5%) |
|  | Sensitivity analysis | 16 (8.3%) |
|  | Stratification or sub-group analyses | 14 (7.3%) |
| Student *t*-test |  | 40 (20.7%) |
|  | One sample *t*-test | 1 (0.5%) |
|  | Paired/matched *t*-test | 0 (0) |
|  | Two independent samples *t*-test | 38 (19.7%) |
|  | Z test | 1 (0.5%) |
| Analysis of variance(ANOVA) |  | 25 (13.0%) |
|  | Completely random design ANOVA | 24 (12.4%) |
|  | Randomized block design ANOVA | 1 (0.5%) |
|  | Factorial design ANOVA | 0 (0) |
|  | Cross-over ANOVA | 0 (0) |
|  | Analysis of covariance | 0 (0) |
|  | Multivariate Analysis Of Variance | 0 (0) |
| Multiple comparisons |  | 5 (2.6%) |
|  | Students-Newman-Keuls method | 0 (0) |
|  | Bonferroni method | 3 (1.6%) |
|  | Dunnett method | 1 (0.5%) |
|  | Duncan's method | 0 (0) |
|  | LSD method | 0 (0) |
|  | Tukey method | 0 (0) |
|  | Sidak method | 0(0) |
|  | Scheffe method | 1 (0.5%) |
|  | FDR (false discovery rate) | 0 (0) |
| Repeated measurement data |  | 12 (6.2%) |
|  | Repeated measurement data ANOVA | 4 (2.1%) |
|  | GEE (Generalized estimating equation) | 2 (1.0%) |
|  | MMRM (Mixed-effect models for repeated measures) | 1 (0.5%) |
|  | GLMM (generalized linear mixed models) | 5 (2.6%) |
| Non-parametric test |  | 44 (22.8%) |
|  | Sign test | 0 (0) |
|  | Wilcoxon signed-rank test | 21 (10.9%) |
|  | Mann-Whitney test | 18 (9.3%) |
| | Kruskal-Wallis $H$ test | 7 (3.6%) |
|  | Friedman test | 0 (0) |
|  | Kolmogorov-Smirnov test | 2(1.0%) |
|  | Median test | 0 (0) |
| Contingency tables |  | 100 (51.8%) |
|  | Chi-square test | 90 (46.6%) |
|  | McNemar's test | 0 (0) |
|  | Fisher's exact test | 31 (16.1%) |
| Correlation analysis |  | 26 (13.5%) |
|  | Pearson correlation coefficient | 5 (2.6%) |
|  | Spearman correlation coefficient | 11 (5.7%) |
|  | Kendall's correlation coefficient | 1 (0.5%) |
|  | Trend test | 12 (6.2%) |
|  | Partial correlation coefficient | 0 (0) |
|  | Multiple correlation coefficient | 0 (0) |
| Multiple regression |  | 106 (54.9%) |
|  | Linear regression | 7 (3.6%) |
|  | Curve estimate | 1 (0.5%) |
|  | Path analysis | 0 (0) |
|  | Logistic regression | 93 (48.2%) |
|  | Poisson regression | 3 (1.6%) |
|  | Negative binominal | 2 (1.0%) |
|  | Spline regression | 0 (0) |
|  | Other regression model | 5 (2.6%) |
| Survival analysis |  | 165 (85.5%) |
|  | Kaplan-Meier estimate | 24(12.4%) |
|  | Life-table method estimate | 1 (0.5%) |
|  | Log-rank test | 10 (5.2%) |
|  | Breslow test | 0 (0) |
|  | Tarone-Ware test | 0 (0) |
|  | Cox proportional hazards model | 41 (21.2%) |
|  | Other survival model | 4 (2.1%) |
| Consistency measurement |  | 2 (1.0%) |
| Principal component analysis |  | 0 (0) |
| Factor analysis |  | 0 (0) |
| Discriminant analysis |  | 0 (0) |
| Cluster analysis |  | 2(1.0%) |
| Log-linear models |  | 0 (0) |
| Structural equation modeling (SEM) |  | 0 (0) |
| Multilevel modeling |  | 0 (0) |
| Multi dimensional scaling analysis |  | 0 (0) |
| Bayesian analyses |  | 0 (0) |
| Other statistical methods |  | 14 (7.3%) |

**Figure. 3.**
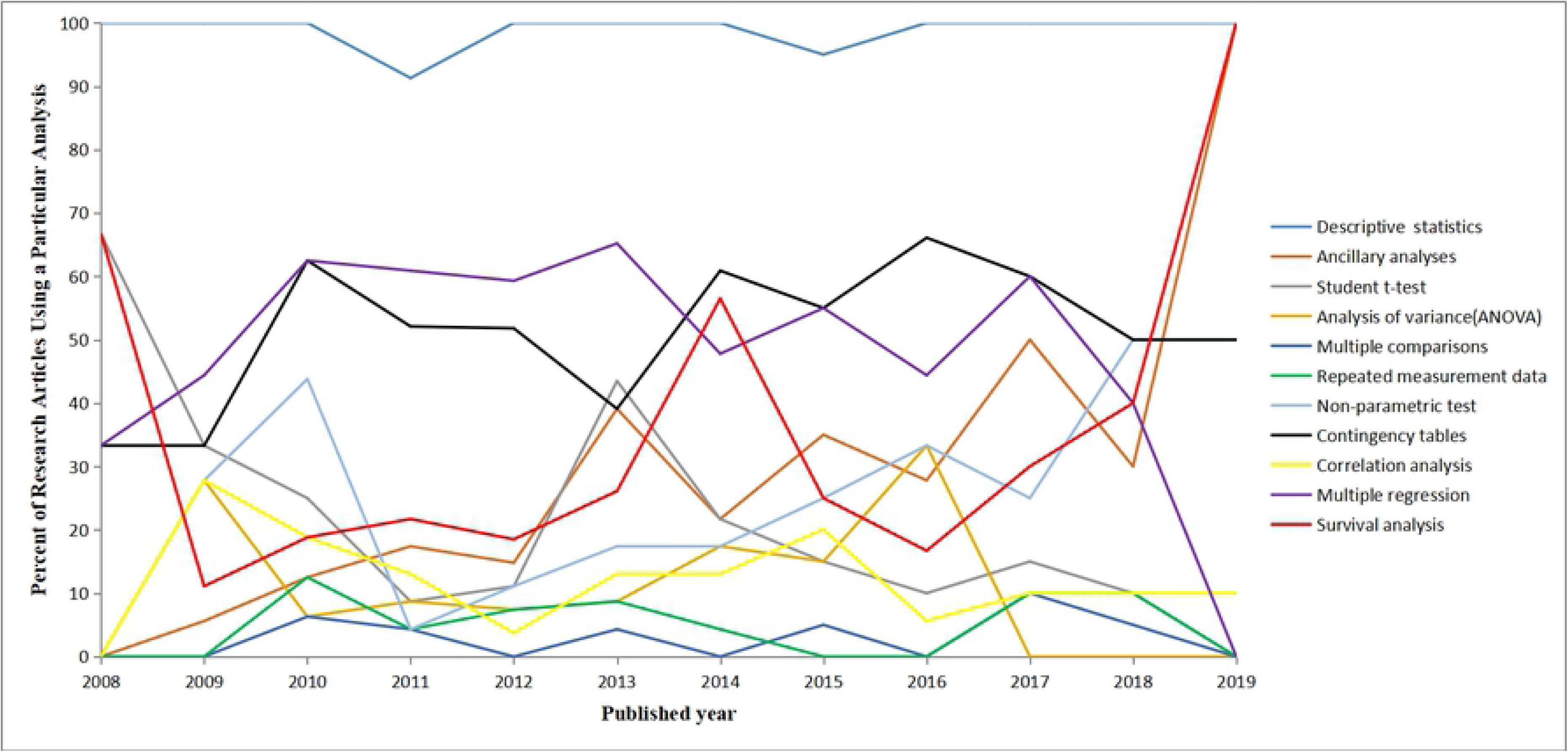
Trends of the statistical methods.

### The reporting quality of statistical methods of included articles

The mean score of reporting quality in included articles was 0.42 (range: 0.11-0.82) with a standard deviation of 0.15. 132 (68.4%) articles have a score less than 0.5. The statistical reporting score has not been improved over the past decade. The articles that number of author greater than or equal to 7 had a higher score than others, but the difference was not statistically significant. The statistical reporting score was similar in three type studies. The articles that have participation of statistician or epidemiologist tend to have a higher statistical reporting score.

As shown in Table.3, three of the seven items that must be reported were often missed: outliers (n=2, 1.0%), sample size (n=13, 6.7%), missing data (n=19, 9.8%). Two items were reported suboptimal: one or two tailed (n=84, 44.9%), confidence intervals (n=134, 69.4%). In addition, 158 articles (84.5%) reported the name of statistical package or program with 161 articles (86.1%) reported exact *P*-values of significance test.

**Table 3.** Reporting of common statistical methods in evaluated journals.

| **Preliminary analysis** |  | Reporting rate |
| --- | --- | --- |
| * | 1. Sample calculation / Power analysis | 6.7% (13/193) |
| * | 2. Whether there were outliers in an article | 1.0% (2/193) |
|  | 3. Report how any missing data were treated | 9.8% (19/193) |
| If mean with standard deviation, median with interpercentile range and proportion were reported in an article, item 4, 5, 6 are considered applicable items. | 4. Mean and standard deviation with examine for normality | 7.8% (154/193) |
|  | 5. Median and interpercentile range with explaining reason (non-normality, ordinal data, quantitative variable without exact data at either end) | 22.8% (44/193) |
|  | 6. Proportion (report numerator and denominator) | 10.9% (21/193) |
| * | 7. Name of statistical package or program reported | 84.5% (158/187<sup>a</sup>) |
| * | 8. Report the alpha level (e.g. 0.05) that defines statistical significance | 65.2% (122/187<sup>a</sup>) |
| * | 9. Report one or two tailed for test. (Justify the use of one-tailed tests.) | 44.9% (84/187<sup>a</sup>) |
| * | 10. Exact P-values for test (e.g. P=0.23) | 86.1% (161/187<sup>a</sup>) |
| * | 11. Confidence intervals | 69.4% (134/193) |
| **Statistical methods** |  |  |
| Student *t*- test |  | n=41 |
|  | 1. Assessing assumptions for *t*-test | 17.1% (7/41) |
|  | 2. Report the type of *t*-test (e.g. one sample, two independent samples, paired/matched) | 70.7% (29/41) |
| ANOVA/ANCOVA |  | n=25 |
|  | 1. Assessing assumptions for ANOVA /ANCOVA | 4.0% (1/25) |
|  | 2. Report the type of ANOVA (e.g. one way, randomized block design, factorial design, repeated measure design, cross-over design etc. ) | 48.0% (12/25) |
| Chi-square test |  | n=96 |
|  | 1. Report the type of Chi-square test (Pearson Chi-square test, continuity correction Chi-square test, McNemar's test or Fisher's exact test) | 74.0% (71/96) |
| Non-parametric test |  | n=39 |
|  | 1. Provide reasons (continuous variable with non-normality, ordinal data, quantitative variable without exact data at either end) | 23.1% (9/39) |
|  | 2. Report the type of non-parametric test (e.g. sign test, Wilcoxon signed-rank test, Mann-Whitney test, Kruskal-Wallis H test, Friedman's test) | 100% (39/39) |
| Linear correlation |  | n=13 |
|  | 1. Report the type of correlation coefficient, with confidence intervals (e.g. pearson, spearman, Kendall's tau_b correlation or trend test) and provide reasons (continuous variable with non-normality, ordinal variable) | 100% (13/13) |
| Multiple linear regression |  | n=9 |
|  | 1. Conformity with a linear gradient | 55.6% (5/9) |
|  | 2. Reason of selection of variables (e.g. based on published literatures, professional knowledge, results of univariate analysis or decision of the researcher) | 22.2% (2/9) |
|  | 3. Methods for variable selection (all possible subsets selection, forward selection, backward selection, stepwise selection) | 55.6% (5/9) |
|  | 4. Interactions among independent variables | 22.2% (2/9) |
|  | 5. Colinearity of independent variables | 11.1% (1/9) |
|  | 6. Coding of variables | 11.1% (1/9) |
|  | 7. Validation of the statistical model | 11.1% (1/9) |
| | 8. Goodness of fit test (e.g. coefficient of determination, $r^2$) | 11.1% (1/9) |
| Multiple Logistic regression |  | n=93 |
|  | 1. Sufficient events (>10) per variable (the ratio of outcome events to independent variables) | 98.9% (92/93) |
|  | 2. Conformity with linear gradient for continuous or rank variables | 26.9% (25/93) |
|  | 3. Reason of selection of variables (e.g. based on published literatures, professional knowledge, results of univariate analysis or decision of the researcher) | 8.6% (8/93) |
|  | 4. Methods for variable selection (forward selection, backward selection, stepwise selection) | 24.7% (23/93) |
|  | 5. Interactions between independent variables | 25.8% (24/93) |
|  | 6. Colinearity of independent variables | 3.2% (3/93) |
|  | 7. Coding of variables | 20.4% (19/93) |
|  | 8. Validation of the statistical model (e.g. likelihood ratio test, Wald test, score test) | 6.5% (6/93) |
|  | 9. Goodness of fit test | 5.4% (5/93) |
| Survival analysis |  | n=49 |
|  | 1. Identify dates or events marking the beginning and the end of the time period analyzed | 28.6% (14/49) |
|  | 2. Report follow-up information (e.g. the mean of follow-up time, the median of follow-up time, the average of follow-up, the range of follow-up, person-years) | 32.7% (16/49) |
|  | 3. Survival rate or survival function | 49.0% (24/49) |
|  | 4. Report the circumstances under which data were censored | 12.2% (6/49) |
| If comparative analysis was applied |  | n=25 |
|  | 1. Report the statistical methods applied to compare two or more survival curves. | 72.0% (18/25) |
|  | 2. Report median survival time | 80.0% (20/25) |
| If Cox model was applied |  | n=42 |
|  | 1. Reported assumption of proportional hazard (assumption of PH) for Cox proportional hazards regression model (Cox model) | 66.7% (28/42) |
|  | 2. Conformity with a linear gradient for continuous or rank variables for Cox model | 31.0% (13/42) |
|  | 3. Interactions between independent variables for Cox model | 40.5% (17/42) |
|  | 4. Colinearity of independent variables for the Cox model | 2.4% (1/42) |
|  | 5. Reason of selection of variables (e.g. based on published literature, professional knowledge, results of univariate analysis or decision of the researcher) for Cox model | 11.9% (5/42) |
|  | 6. Methods for variable selection (forward selection, backward selection, stepwise selection) for Cox model | 16.7% (7/42) |
|  | 7. Coding of variables for Cox model | 16.7% (7/42) |

The quality of the multiple analyses reporting left much to be desired. The mean of reporting score of the multiple logistic regression was 0.24, 0.27 for multiple cox regression, and 0.29 for multiple linear regression. The item that was reported most often in multiple logistic regression was ‘‘sufficient events’’ (n=92, 98.9%) followed by conformity with linear gradient (n=25, 26.9%) and interactions between independent variables (n=24, 25.8%). The least-reported item was colinearity analysis (n=3, 3.2%), and the goodness of fit test was slightly better (n=5, 5.4%). Methods for variable selection (n=23), coding of variables (n=19), reason of selection of variables (n=8) and validation of the statistical model (n=6) were reported in 24.7%, 20.4%, 8.6% and 6.5% of the articles, respectively. Of the reporting of Cox regression models, assumption of proportional hazard were reported in 66.7% studies (n=28). However, important information such as interaction test (n=17, 40.5%), methods for variable selection (n=7, 16.7%), variables coding (n=7, 16.7%), reason of selection of variables (n=5, 11.9%) and colinearity test (n=1, 2.4%) were rarely reported. Seven articles (17.1%) assessed assumptions for t-test and 1 article for ANOVA (4.0%). Nine articles (23.1%) described the reason for conducting non-parametric tests.

The included articles were divided into inferior (n=144) and superior reporting quality groups (n=49) based on the cut-off value (0.50). As shown in table.4, univariate logistic regression analyses showed the following factors were related to the high reporting quality: participation of a statistician or epidemiologist (OR=1.92, 95%CI=1.04-3.71) and number of authors (OR=2.23, 95%CI=1.13-4.37).

Multivariate logistic regression analysis likewise demonstrated that participation of a statistician or epidemiologist (OR=1.73, 95%CI=1.18-3.39) and number of authors (OR=2.06, 95%CI=1.04-4.08) were associated with the superior reporting quality. The multivariate logistic model has sufficient events (the ratio of outcome events to independent variables was 24.5). The likelihood ratio test was used to validate the goodness of fit of the model, it was found that the fitting degree of the model was good (χ^2^=0.84, *P*=0.656). The max variance inflation factor (VIF) was 1.026 to indicate no multicollinearity among the independent variables.

**Table 4.**
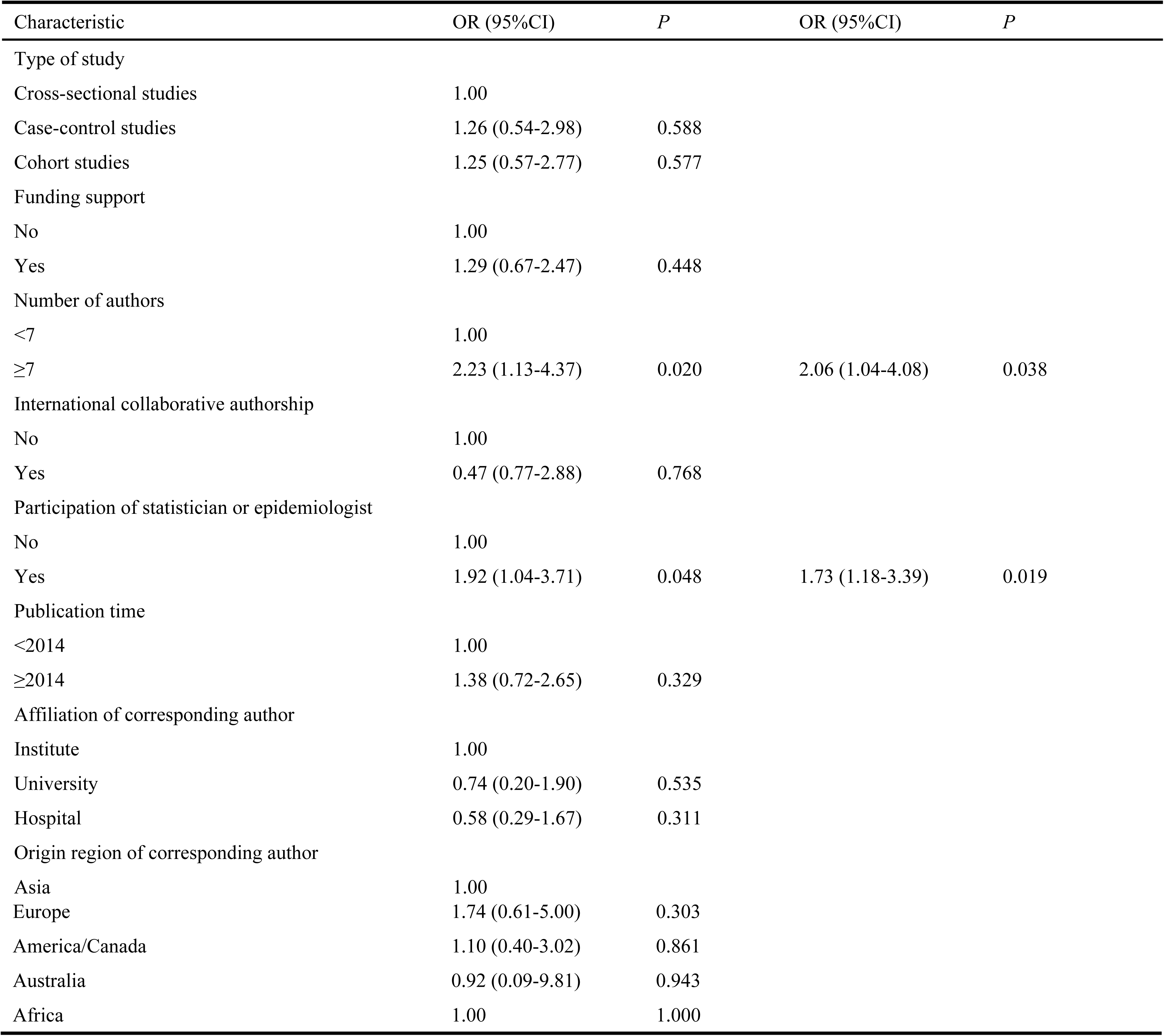
Univariate and multivariate logistic regression analyses of predictive factors associated with superior statistical methods reporting quality

## Discussion

The study estimated the frequency, trend and reporting quality of statistical methods of 193 observational studies published in five high-impact urology journals after 2008. The number and complexity of statistical methods have not changed in recent years, and the reporting quality of statistical methods has not improved either.

Compared with other disciplines in the medical field, fewer statistical methods were used in urology studies (15, 16). This is an alarming phenomenon, because researchers need to use several different statistical methods when facing different distribution data or different study purposes. Logistic regression models and chi-square test are most frequently used statistical methods in urological observational studies, because observational studies make heavy use of binary endpoints. Different from previous research results, our study indicated that the statistical methods of observational studies in urology have not changed more complicated in the past years (15-17). During recent years, statisticians have introduced new and complex methods that are attributable to the rapid expansion in computing capability, such as, Bayesian methods, artificial neural networks, and machine learning. However, we found no reference to these methods in any of the evaluated articles. These may be on account of the new complex statistical methods are not included in the introductory or second-level statistics courses, few authors are familiar with these methods. 36 (18.7%) articles only used simple tests, however, the use of simple statistical methods only, may be generally insufficient. Sophisticated and valid methods should be used based on study designs and data properties to avoid the bias and inflation of the type I error rate for the treatment effect.

Standardized statistical reporting of medical papers not only help editors or reviewers better understand research design to improve the quality of journal papers, but also enable readers in related fields to better understand the content and results of research to enrich their professional knowledge. However, the reporting quality of statistical methods in this study was inferior and still needs to be improved in many aspects. The average score of reporting quality in included articles was 0.42, and 68.4% articles have a score less than 0.5, this means that the statistical reporting adherence of most articles was less than 50%.

Pre-study calculation of the sample size is necessary. The correct sample size of a study has the advantage of enhancing feasibility, reducing costs, and also has ethical implications. If the sample size is too small, it can not detect the effect, causing type **II** errors. Too large sample size may not only waste time, resources and money, but also difficult to implement. The Strengthening the Reporting of Observation Studies in Epidemiology (STROBE) statement also recommends researchers should explain how the study size was arrived at when they conducting observational studies (18, 19). However, only 6.7% articles reported the calculation of sample size or power analysis in this study. Data missing is a common and unavoidable problem in the medical studies. If the missing data are not handled properly, it will cause bias or insufficient use of data, thus reducing the efficiency of the study or biased inferences (20, 21). Nevertheless, we found that only 9.8% included studies indicated number of variables with missing data. Multivariate analyses are very sensitive to outliers, as outliers may cover or cause multicollinearity between independent variables and affect the parametric estimation of the model. However, just 1% articles detected outliers of the original data. The encouraging finding in this study was that the statistical software (84.5%) and exact *P* values (86.1%) were reported at high rates as essential requirements for statistical analysis. But there is still room for improvement. In statistical inference, it is advisable to report the exact *P*-values and confidence intervals. A large confidence interval means that the small sample size and large random error. In this way, the conclusion drawn by the exact *P* is questionable.

We found that 133 articles (68.9%) employed simple statistical methods and several problems were occurred in the reporting of these methods. The use of the t test and ANOVA requires that the data follow normal distribution and homogeneity of variance, whereas very fewer articles mention it. Using non-parametric tests on data suitable for parametric tests will reduce statistical power. Therefore, the reason for using non-parametric tests should be stated. However, only 23.1% articles described the reasons.

The confounders controlling is a crucial step in analytical observational studies, and multivariable analyses are widely used as statistical adjustment techniques (22). 151 articles (78.2%) applied multivariable regression models such as logistic, Cox, linear, or Poisson regression. Several previous studies have shown that there was still room for improvement in quality of the reporting of multivariate regression (13, 22-24), which was consistent with our findings. Only 6 articles (5.9%) described goodness of fit test. By contrast, Casals *et al*. conducted a systematic review of 108 articles, and found testing for goodness-of-fit was reported in 15.7% (23). 96.5% included articles did not consider collinearity diagnostics, which may result unstable regression coefficients or wide confidence intervals, and even affect the selection of variables. Real, J *et al.* found that 26.2% articles that used multivariable analyses described linear gradient for continuous or rank variables (22). This reporting rate in our study was 40.3%, higher than their findings, but far from ideal as well. This may be due to the lack of automatic options for this test in current common statistical software.

The result of multivariate logistic regression analysis demonstrated that the participation of a statistician or epidemiologist was associated with high reporting quality. Professional statisticians or epidemiologists can not only provide correct guidance for data processing in research, but also better understand what important information in statistical analysis should be reported. We recommend that researchers seek the assistance of statistical professionals when processing data and writing articles.

Some limitations were present in our study as well. First, we only searched five top journals and not the full breadth of the articles related to urology. However, the highest impact factor journals have the utmost visibility in the urology literature and are likely more relied upon by doctors to inform practice. Second, only the most commonly used statistical methods are included in our checklists, and some rarely used statistical methods are not included.

## Conclusion

The statistical reporting of published observational studies in 5 high-impact factor urological journals was often ambiguous, especially for multivariable regression models, and there is room for improvement. The participation of statistician or epidemiologist may improve statistical reporting quality. The authors, reviewers and editors should increase their knowledge of statistical methods, especially new and complex methods. We recommend editorial board of journal publish guidelines to guide and facilitate critical appraisal of statistical reporting.

## Supporting Information

**Appendix S1** Common statistical methods in medical studies (Docx)

**Appendix S2** Reporting of common statistical methods in evaluated journals (Doc)

## Author contribution

Conceived and designed the experiments: DSY ZXB LBB. Performed the experiments: DSY XH ZJG. Analyzed the data: DSY LBB. Contributed reagents/materials/analysis tools: DSY ZXB LBB. Contributed to the writing of the manuscript: DSY LBB ZXB.

## References

1. Fosang AJ, Colbran RJ. Transparency Is the Key to Quality. The Journal of biological chemistry. 2015;290(50):29692–4.

2. Nature. Reducing our irreproducibility. Nature. 2013;496:398(DOI: https://doi.org/10.1038/496398a).

3. Wu R, Glen P, Ramsay T, Martel G. Reporting quality of statistical methods in surgical observational studies: protocol for systematic review. Systematic reviews. 2014;3:70.

4. Schor S, Karten I. Statistical evaluation of medical journal manuscripts. Jama. 1966;195(13):1123–8.

5. Yoon U, Knobloch K. Quality of reporting in sports injury prevention abstracts according to the CONSORT and STROBE criteria: an analysis of the World Congress of Sports Injury Prevention in 2005 and 2008. Br J Sports Med. 2012;46(3):202–6.

6. Lang TA, Altman DG. Basic statistical reporting for articles published in biomedical journals: the “Statistical Analyses and Methods in the Published Literature” or the SAMPL Guidelines. International journal of nursing studies. 2015;52(1):5–9.

7. Pentti Nieminen JIV. An instrument to assess the statistical intensity of medical research papers. PloS one. 2017.

8. Nieminen P, Toljamo T, Vahanikkila H. Reporting data analysis methods in high-impact respiratory journals. ERJ open research. 2018;4(2).

9. Emerson JD, Colditz GA. Use of statistical analysis in the New England Journal of Medicine. The New England journal of medicine. 1983;309(12):709–13.

10. Scales CD, Norris RD, Peterson BL, Preminger GM, Dahm P. Clinical Research and Statistical Methods in the Urology Literature. Journal of Urology. 2005;174(4 Part 1):1374–9.

11. Guglielminotti J, Dechartres A, Mentre F, Montravers P, Longrois D, Laouenan C. Reporting and Methodology of Multivariable Analyses in Prognostic Observational Studies Published in 4 Anesthesiology Journals: A Methodological Descriptive Review. Anesthesia and analgesia. 2015;121(4):1011–29.

12. Weissgerber TL, Garcia-Valencia O. Why we need to report more than ‘Data were Analyzed by t-tests or ANOVA’. 2018;7.

13. Zhang YY, Zhou XB, Wang QZ, Zhu XY. Quality of reporting of multivariable logistic regression models in Chinese clinical medical journals. Medicine. 2017;96(21):e6972.

14. Zhu X, Zhou X, Zhang Y, Sun X, Liu H, Zhang Y. Reporting and methodological quality of survival analysis in articles published in Chinese oncology journals. Medicine. 2017;96(50):e9204.

15. Gosho M, Sato Y, Nagashima K, Takahashi S. Trends in study design and the statistical methods employed in a leading general medicine journal. Journal of clinical pharmacy and therapeutics. 2018;43(1):36–44.

16. Gosho M, Nagashima K, Takahashi S, Ware JH, Laird NM. Statistical Methods in the Journal - An Update. The New England journal of medicine. 2017;376(11):1086–7.

17. Horton NJ, Switzer SS. Statistical methods in the journal. The New England journal of medicine. 2005;353(18):1977–9.

18. von Elm E, Altman DG, Egger M, Pocock SJ, Gotzsche PC, Vandenbroucke JP. Strengthening the Reporting of Observational Studies in Epidemiology (STROBE) statement: guidelines for reporting observational studies. BMJ (Clinical research ed). 2007;335(7624):806–8.

19. Vandenbroucke JP. The making of STROBE. Epidemiology. 2007;18(6):797–9.

20. Zhang Z. Missing data exploration: highlighting graphical presentation of missing pattern. Annals of translational medicine. 2015;3(22):356.

21. Tseng CH, Chen YH. Regularized approach for data missing not at random. Statistical methods in medical research. 2019;28(1):134–50.

22. Real J, Forne C, Roso-Llorach A, Martinez-Sanchez JM. Quality Reporting of Multivariable Regression Models in Observational Studies: Review of a Representative Sample of Articles Published in Biomedical Journals. Medicine. 2016;95(20):e3653.

23. Casals M, Girabent-Farres M, Carrasco JL. Methodological quality and reporting of generalized linear mixed models in clinical medicine (2000-2012): a systematic review. PloS one. 2014;9(11):e112653.

24. Kumar R, Indrayan A, Chhabra P. Reporting quality of multivariable logistic regression in selected Indian medical journals. Journal of postgraduate medicine. 2012;58(2):123–6.

